# Distinct mechanisms of CB1 and GABA_B_ receptor presynaptic modulation of striatal indirect pathway projections to mouse Globus Pallidus

**DOI:** 10.1101/2022.07.21.500979

**Authors:** Giacomo Sitzia, Karina Possa Abrahao, Daniel Liput, Gian Marco Calandra, David M. Lovinger

## Abstract

Presynaptic modulation is a fundamental process regulating synaptic transmission. Striatal indirect pathway projections originate from A2A- expressing spiny projection neurons (iSPNs), targeting the globus pallidus external segment (GPe) and control the firing of the tonically active GPe neurons via GABA release. It is unclear if and how the presynaptic GPCRs, GABA_B_ and CB1 receptors, modulate iSPN-GPe projections. Here we used an optogenetic platform to study presynaptic Ca^2+^ and GABAergic transmission at iSPN projections, using a genetic strategy to express the calcium sensor GCaMP6f or the excitatory channelrhodopsin (hChR2) on iSPNs. We found that P/Q-type calcium channels are the primary VGCC-subtype controlling presynaptic calcium and GABA release at iSPN-GPe projections. N-type and L-type VGCCs contribute to GABA release at iSPN-GPe synapses. GABA_B_ receptor activation resulted in a reversible inhibition of presynaptic Ca^2+^ transients (PreCaTs) and an inhibition of GABAergic transmission at iSPN-GPe synapses. CB1 receptor activation did not inhibit PreCaTs while inhibiting GABAergic transmission at iSPN-GPe projections. CB1 effects on GABAergic transmission persisted in experiments where Na_V_ and K_V_1 were blocked, indicating a VGCC- and K_V_1 independent presynaptic mechanism of action of CB1 receptors. Taken together, presynaptic modulation of iSPN-GPe projections by CB1 and GABA_B_ receptors is mediated by distinct mechanisms.

**Key Points:**

P/Q-type are the predominant VGCC controlling presynaptic Ca^2+^ and GABA release on the striatal indirect pathway projections
GABA_B_ receptor modulate of iSPN-GPe projections via a VGCC- dependent mechanism
CB1 receptors modulate iSPN-GPe projections via a VGCC- independent mechanism

## Introduction

The dorsal striatum (DS) is the main input structure of the basal ganglia (BG), a group of subcortical regions involved in motor control, associative learning and decision making. Spiny projection neurons (SPNs) constitute up to 95% of neurons in the rodent striatum (Graveland & Di Figlia, 1985). SPNs are classified based on their molecular markers and projection targets into direct pathway SPNs (dSPNs) and indirect pathway SPNs (iSPNs) (Gerfen et al., 1990). The dSPNs express the dopamine D1 receptor and project to the substantia nigra pars reticulata (SNr), the entopeduncular nucleus (EPN) and, through axonal collaterals, to the globus pallidus external segment (GPe) (Chang et al., 1981). iSPNs express dopamine D2 and adenosine A2A receptors and project to the GPe. Neurons in the GPe are tonically active, and this activity is influenced by GABAergic inputs from the iSPNs. Presynaptic neuromodulation by GPCRs including GABA_B_ and CB1 controls GABA release in the GPe (Engler et al., 2006; Kaneda & Kita, 2005) but the synapse- specific effects and mechanisms of action are unclear. Thus, understanding how GPCRs govern presynaptic calcium entry and neurotransmitter release at the iSPN projections to the GPe will allow to better determine how the striatum controls its downstream targets, in physiology and in disease.

Two key modulators of GABAergic transmission in the GPe are the G_i/o_ coupled cannabinoid type 1 (CB1) and GABA_B_ receptors. Expression levels of CB1 receptors on striatal terminals in the GPe and SNr are among the highest in the CNS (Davis et al., 2018; Hu & Mackie K., 2015). The expression levels of CB1 receptors in the GPe have been shown to drop dramatically when striatopallidal projections are lesioned (Hernkenham et al., 1991) or if CB1 receptors are genetically knocked out from SPNs (Bonm et al., 2021). To date, it remains unclear whether pallidal neurons express functional CB1 receptors. CB1 receptors control GABA release in the GPe (Engler et al., 2006) but it is unknown what set of GABAergic synapses mediates this effect. Sources of extra-striatal GABAergic inputs in the GPe include local collaterals (Ketzef et al., 2021; Higgs et al., 2021) and GPi inputs (Weglage et al., 2022). GABA_B_ receptors are expressed on striatopallidal projections (Chen et al, 2004). GABA_B_ receptors exert pre-synaptic effects by reducing GABA release at GABAergic synapses in the GPe but also act postsynaptically to inhibit GPe neurons (Kaneda & Kita, 2005). Overall, it remains unclear if and how CB1 and GABA_B_ receptors modulate transmission specifically at iSPN-GPe projections.

CB1 and GABA_B_ receptors suppress neurotransmitter release in the CNS through signaling cascades mediated by the G_βγ_ subunit, including inhibition of voltage-gated calcium channels (VGCCs), inhibition of the vesicle release complex downstream of VGCCs and activation of K^+^channels (Araque et al., 2017; Chalifoux & Carter, 2011). Mechanisms involving the G_αi_/G_o_ subunits, including the inhibition of cAMP/ PKA signaling, also influence neurotransmitter release (Araque et al., 2017; Chalifoux & Carter, 2011). The downstream mechanisms engaged by CB1 and GABA_B_ receptors are synapse specific and can be non- canonic. In the nucleus accumbens (NAc), CB1 receptors suppress neurotransmitter release at cortico-accumbal synapses through a VGCC-independent, K^+^ channel-dependent mechanism (Robbe et al., 2001), while GABA_B_ receptors inhibit neurotransmitter release at glutamatergic synapses through a VGCC-independent, SNAP25-dependent mechanism (Manz et al., 2019). In the DS, GABA_B_ receptors suppress neurotransmitter release at corticostriatal synapses via a VGCC-dependent mechanism (Kupfershmidt & Lovinger, 2015).

Here we studied the presynaptic mechanisms of action of CB1 and GABA_B_ receptors on the iSPN-GPe projections. We validated a slice photometry approach to measure presynaptic Ca^2+^ at iSPN-GPe projections, using transgenic mice expressing the Ca^2+^ indicator GCaMP6f in iSPNs (A2aCre-GCaMP6f mice). We also used transgenic mice expressing the excitatory opsin channelrhodopsin (hChR2) to study the mechanisms regulating CB1 and GABA_B_ receptors effects on GABAergic transmission at iSPN-GPe synapses. We found that P/Q-type VGCCs are the primary VGCCs mediating Presynaptic Calcium Transients (PreCaTs) and GABAergic transmission at iSPN synapses in the GPe. N- and L-type VGCCS inhibition reduced GABAergic transmission, although with a smaller effect size than P/Q-type VGCCs, but not PreCaTs. GABA_B_ receptor activation decreased PreCaTs and GABAergic transmission. Conversely, CB1 receptor activation at iSPN projections did not decrease PreCaTs, while reducing GABAergic transmission. CB1 effects persisted when potassium voltage-gated channels 1 (K_v_1) and sodium voltage gated channels (Na_V_) were blocked, and reduced the frequency of miniature inhibitory postsynaptic currents (mIPSCs), which we found to be insensitive to the VGCC blocker cadmium in the GPe. Taken together, our results indicate that CB1 receptor inhibition of GABA release at iSPN terminals is mediated by a K_V_1, Na_V_ and VGCC-independent mechanism.

## Methods

### Ethical approval

Experiments were conducted in accordance with the National Institutes of Health’s Guidelines for Animal Care and Use, and all experimental procedures were approved by the National Institute on Alcohol Abuse and Alcoholism Animal Care and Use Committee (Protocol LIN-DL-1).

### Animals

Mice were housed in groups of 2-4 in the NIAAA vivarium on a 12 hr light cycle with ad libitum access to food and water. B6;129S-Gt(ROSA)26Sortm95.1(CAG-GCaMP6f)Hze/J (GCaMP6f) and B6.Cg-Gt(ROSA)26Sortm32(CAG-COP4*H134R/EGFP)Hze/J (Ai14-ChR2) mice were purchased from The Jackson Laboratory (JAX; Bar Harbor, ME, USA). B6.FVB(Cg)-Tg(Adora2a-cre)KG139Gsat/Mmucd (A2A-Cre) mice were purchased from the Mutant mouse resource and research centers (MMRRC, 030869, Davis, CA, USA). These mutant mouse lines were maintained by breeding with wildtype C57BL6J mice purchased from JAX. Experimental animals were generated by breeding male or female mice homozygous for the GCaMP6f or ChR2 allele with mice heterozygous for the A2A-Cre alleles to restrict GCaMP6f or ChR2H134R (hChR2) expression to striatal indirect pathway SPNs. Male and female mice (aged 3-24 weeks) were included in the electrophysiology and slice photometry experiments experiments. No differences were observed in effect sizes of the drugs used across different ages. Male mice were used to validate GFP expression in GCaMP6f mice (aged 70w). Genotyping was performed by polymerase chain reaction (PCR) on genomic DNA from ear biopsies.

### Immunohistochemistry and confocal imaging

A2aGCaMP6 f mice (n = 3), were anesthetized with pentobarbital (50 mg/ kg) prior to the transcardiac perfusion of 1× PBS, followed by 4% formaldehyde (FA) (4% paraformaldehyde in 1× PBS). Brains were kept in 4% FA overnight and then transferred to 1x PBS until slicing. Free-floating sections (50 mM) were cut using a Pelco easySlicer Vibratome (Ted Pella Inc., Redding, Ca, USA). Slices were washed 3 × 5 minutes in 1× PBS, and then incubated in a blocking solution containing 5% normal goat serum in PBS-T (0.2% Triton X-100). Slices were then incubated for 48 hr at 4C in 1× PBS containing the primary antibodies (Chicken anti-GFP, AbCam cat. No. #13970. 1:2000; Rabbit anti-NeuN, Cell Signaling Technology, cat. No. #24307, 1:1000). Slices were then washed 3 × 5 minutes in 1x PBS and incubated in 1x PBS containing the secondary antibodies for 2 hours (Alexa Fluor 488, Goat-anti-chicken, Invitrogen A11039, 1:500; Alexa Fluor 594 goat anti-rabbit, Invitrogen A11012, 1:500). A final wash of 3 × 5 minutes was performed prior to mounting using DAPI Fluoromount-G^®^ (Southern Biotech, Cat No. 0100-20). Slices were imaged on a Zeiss LSM 800 laser confocal scan head mounted on a Zeiss Z1Axio Observer inverted microsope frame (Carl Zeiss, Oberkochen, Germany). An argon laser was used for 488 excitation (Alexa Fluor 488, dichroic MBS 488/594, emission 497-590), a HeNe594 laser for 594 excitation (Alexa Fluor 594, dichroic MBS 488/561, emission 598-670). Objectives used for imaging included a C-Apochromat 63×/1.2 NA W Korr M27 (Zeiss) and Plan-Apochromat 20×/0.8 NA.

### Brain slice preparation for slice photometry and patch-clamp experiments

Mice were anesthetized with isoflurane and decapitated after assessing deep anesthesia by tail-pinch. The brain was rapidly collected and transferred to a slicing chamber filled with ice-cold sucrose-based cutting solution containing the following (in mM): 194 sucrose, 30 NaCl, 4.5 KCl, 26 NaHCO_3_, 1.2 NaH_2_PO_4_, 10 D -glucose, 1 MgCl_2_, saturated with 95% O_2_/5% CO_2_. Coronal sections (250-300 μm) containing the GPe were obtained using a Leica VT1200S Vibratome (Leica Microsystems, Buffalo Grove, IL) and rapidly moved to an incubation chamber filled with artificial cerebrospinal fluid (aCSF) containing (in mM): 124 NaCl, 4.5 KCl, 26 NaHCO_3_, 1.2 NaH_2_PO_4_, 10 D-glucose, 1 MgCl_2_, and 2 CaCl_2_, saturated with 95% O_2_/5% CO_2_. Slices were incubated for 30-60 minutes at 32°C and then maintained at room temperature prior to recording. The same slicing protocol was used for slice photometry and patch-clamp experiments.

### Slice photometry

Photometry recordings were acquired using a Zeiss upright epifluorescence microscope equipped with a 40x/0.8 NA water immersion objective. Slices were placed in a recording chamber and superfused at ~2 mL min-1 with aCSF maintained at ~30°C. GCaMP6f was excited using either a mercury HBO lamp, gated with a uniblitz shutter (Vincent Associates, Rochester, NY, USA) or a 470 nm LED (ThorLabs, Newton, NJ, USA) and filtered at 470 ± 20 nm. Excitation power was measured at the sample plane using a microscope slide photodiode power sensor (ThorLabs) and was 3.8 mW for the mercury HBO lamp and <1.0 mW for the 470 nm LED. Emitted light was collected and filtered at 525 ± 25 nm, passed through a 180 μm2 diaphragm and directed to a photomultiplier tube (PMT; Model D-104, Photon Technology International, Edison, NJ, USA). The PMT output was filtered at 1 kHz and digitized at 6.67 kHz using a Digidata 1322A and Clampex pClamp 10.6 software (Axon Instruments, Molecular Devices LLC, Sunnyvale, CA, USA). A bipolar stimulating electrode was placed in the lateral GPe, just inside the border between the striatum and GPe, and the 180 μm^2^ field of view was positioned just lateral to the electrode tip. Photometry sweeps were acquired every 30 seconds. During each sweep, after an initial rapid decay in fluorescence, a presynaptic calcium transient was evoked by a train of 4 electrical pulses at 10Hz. Slices were equilibrated to the recording chamber conditions and electrical stimulation for ~20 min to allow for response stabilization. Photometry sweeps were averaged over 2 minutes (4 sweeps) before calculating peak ΔF/F values for electrically evoked presynaptic calcium transients (PreCaTs). For experiments using the mercury HBO lamp, the baseline fluorescence intensity tended to decay throughout the entire sweep so a baseline correction procedure was employed as described previously (Kupferschmidt and Lovinger, 2015). However, for experiments using the LED, the baseline fluorescence intensity tended to stabilize prior to evoking presynaptic calcium transients so the baseline correction procedure was omitted.

### Whole-cell patch-clamp recordings

Hemislices were transferred to a recording chamber perfused with aCSF at 30–32 °C with a ~2 ml/min flow rate. Neurons were visualized with an upright microscope (Scientifica, Uckfield, East Sussex, UK) using a LUMPlanFL N × 40/0.80 NA W objective (Olympus, Waltham, MA). Recordings from GPe neurons were obtained using micropipettes (2–4MΩ) made from 1.5 mm OD/1.12 mm ID borosilicate glass with a filament (World Precision Instruments, Sarasota, FL) pulled with a P-97 puller (Sutter Instruments Novato, CA).

Recordings were performed using a Multiclamp 700A amplifier and a Digidata 1322A or Multiclamp 700B and a digidata 1550B digitizer using a low-pass filter of 2 kHz and a sampling frequency of 10 kHz. Data were analyzed using pClamp 10.3 or pClamp 10.6 software (Molecular Devices, Sunnyvale, CA). Most experiments were performed filling micropipettes with a high-chloride solution containing (in mM): 150 CsCl, 10 HEPES, 2.0 MgCl_2_, 0.3 Na-GTP, 3.0 Mg-ATP, 0.2 BAPTA-4Cs, 5 QX-314. Validation of neurotransmitter release at iSPN-GPe projections (Fig. 1) was performed filling micropipettes with a low-chloride solution containing (in mM): 114 CsMeSO3, 5.0 NaCl, 1.0 TEA-Cl, 10 HEPES, 5.0 QX-314, 1.1 EGTA, 0.3 Na-GTP, 4.0 Mg-ATP. Recordings were obtained in voltage clamp mode and neurons were clamped at −45 mV unless specified differently (Fig. 1). GABAA mediated spontaneous (sIPSCs) and optically evoked (oIPSCs) were isolated through the bath-perfusion of aCSF containing the AMPA receptor blocker DNQX (10 μM) and NMDA receptor blocker DL-AP5 (50 μM) and abolished by Gabazine application (Fig. 1). To monitor series resistance (range <30 MΩ) throughout the recording, a 10mV, 50-100 ms depolarizing step was applied every minute; cells were excluded if series resistance varied >20% during the recording.

**Figure 1.**
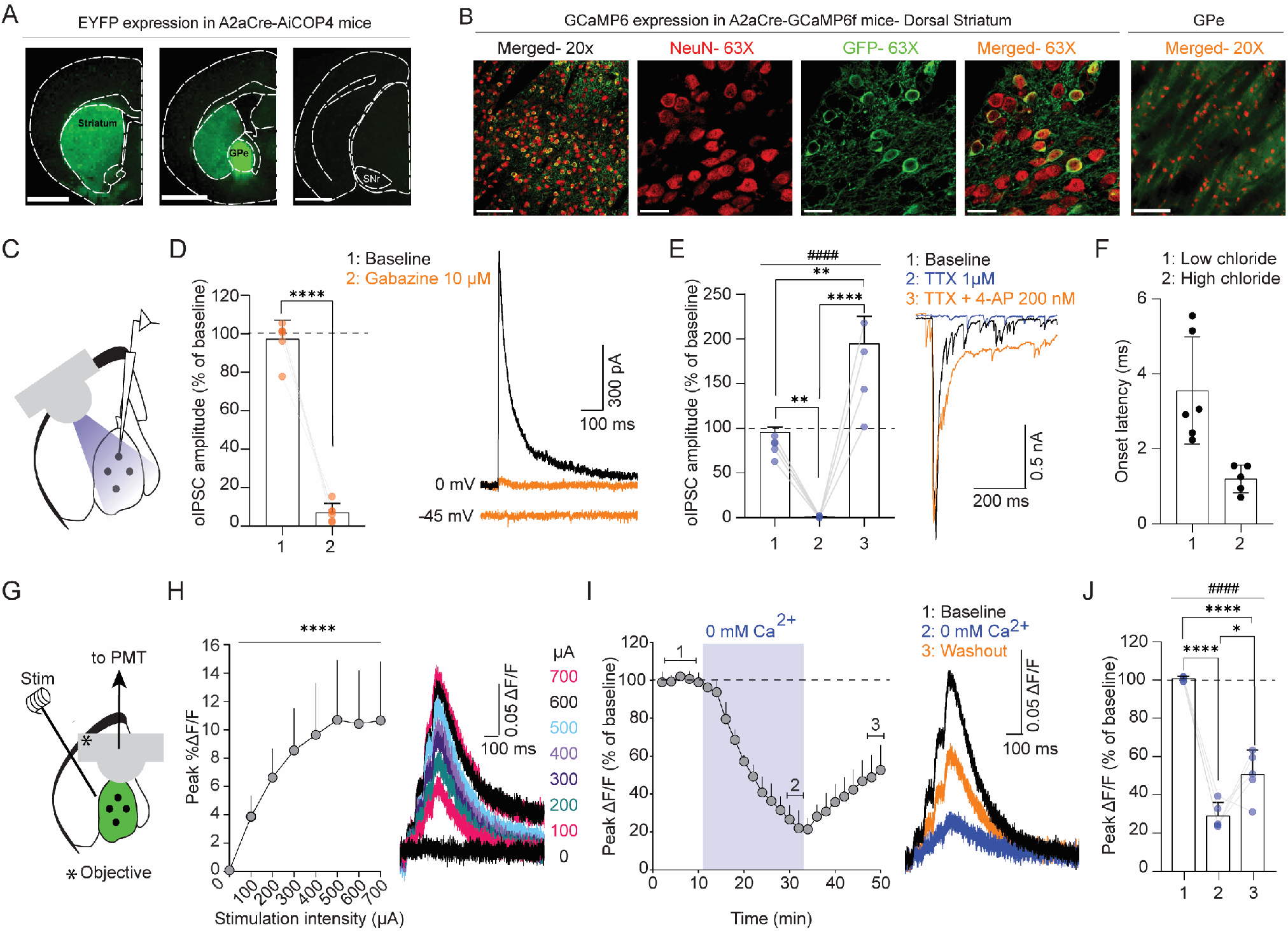
An optogenetic strategy to study GABAergic transmission and PreCaTs at iSPN-GPe projections. **A**: Representative micrographs of EYFP expression in the A2aCre-AiCOP4 line. From left to right: coronal slices containing the dorsal striatum, GPe and SNr. Dense EYFP fluorescence was observed in the striatum and GPe, not in the SNr. **B**: Representative confocal micrographs where IHC staining was performed in A2aCre-GCaMP6f mice using anti-GFP (green) and anti-NeuN (red). From left to right: 20× images of dorsal striatum (scale bar: 100 μM); 63× images of NeuN, GFP and the two channels merged in the dorsal striatum (scale bar: 25μM); 20x image of GPe (scale bar: 100 μM). **C:** Diagram of optogenetic and whole-cell patch clamp experiments. **D**: Summary bar graph and example traces of oIPSC inhibition by Gabazine. Gabazine fully inhibited oIPSCs recorded at iSPN-GPe synapses indicating their GABA_A_ dependency (n = 6 cells from n = 3 mice). **E**: Summary bar graph and example traces of whole-cell patch clamp experiments where (1) baseline, (2) TTX effect and (3) TTX + 4-AP effects were compared. TTX fully inhibited oIPSCs, which were restored above baseline levels by TTX+ 4-AP application (n = 5 cells from n = 4 mice). **F**: onset latency of oIPSCs recorded using high chloride or low chloride intracellular solutions, cells from panels D and E. **G**: schematic of slice photometry experiments. **H**: Input/output (I/O) curve experiment. The amplitude of the PreCaTs scaled with stimulation intensity indicating activity- dependence (n = 5 slices, n = 4 mice). **I:** Time course of Ca^2+^ free ACSF application effects on PreCaTs. **G:** Bar graph summary of effects: baseline, last 2 minutes of Ca^2+^ free ACSF, last 2 minutes of washout (n = 5 slices from n = 4 mice). * P < 0.05; ** P < 0.001; *** P < 0.0001 ****/ #### P < 0.0001.

Optogenetic stimulation protocols: Blue light (470 nm) was delivered through a 4-channel LED driver (ThorLabs, MD, USA). In most experiments oIPSCs were evoked using a train of 4 pulses delivered at 10 Hz. Prior to the start of the recording the light intensity (range 0.4- 6 mW) or the pulse width (0.4-2 ms) were adjusted to obtain a stable oIPSC. For the electrophysiological experiments presented in Fig. 1 and the experiment in fig. 4 where oIPSCs were recorded in presence of TTX+ 4-AP, a 5 ms stimulus with a stimulation intensity of ~10 mW was used to ensure maximal stimulation of presynaptic terminals. As indicated in Fig. 1, oIPSCs were TTX-sensitive and Gabazine sensitive.

### Drugs

Stocks of ω-conotoxin GVIA, ω-agatoxin IVA, (R)-Baclofen, DNQX disodium salt, TTX, 4-AP and DL-AP5 sodium salt were dissolved in ddH2O. WIN55,212-2 mesylate and nifedipine were dissolved in 100% DMSO. All drug stocks were stored at −20°C and diluted in aCSF just prior to superfusion. QX-314 bromide was dissolved fresh in internal solution. ω-conotoxin GVIA and ω-agatoxin IVA were purchased form Alomone Labs (Jerusalem, Israel), QX-314 bromide and 4-AP from Millipore Sigma (St. Louis, MO, USA), and all other drugs from Tocris Biosciences (Minneapolis, MN, USA).

### Statistical Analysis

Statistical analysis was performed in GraphPad Prism (GraphPad Software, La Jolla, California) and Excel. Significance was set at p < 0.05 in all analyses. All data are reported as mean + SD. Exact P values for all comparisons are reported in the main text and all figure legends. Statistical significance is indicated in the figures and figure legends. For all photometry and electrophysiology experiments, except for sIPSC and mIPSC experiments, the data are expressed as % of baseline (defined as the average of the last 8 - 10 minutes recorded prior to drug application). One-way rmANOVA or mixed-effect analysis with no correction followed by Tukey’s multiple comparisons test was performed in experiments where 3 time points were compared (baseline: average of minute 3-8 of baseline; drug: last 2 minutes of drug application, washout: last 2 minutes of washout). Two-way rmANOVA with no correction followed by Tukey’s multiple comparisons test was performed to compare the effects of application of different VGCC blockers on PreCaTs and oIPSCs. Experiments where 2 time points (baseline, drug) were compared were analyzed using paired t-test, if the difference between pairs followed a normal distribution assessed through the Shapiro-Wilk test, or Wilcoxon matched pairs test when not normally distributed.

## Results

### An optogenetic strategy to study presynaptic calcium transients and GABAergic transmission at indirect pathway projections to the GPe

To study presynaptic calcium and GABAergic transmission at iSPN-GPe projections we crossed Adora2a -Cre (A2a-Cre) mice with either GCaMP6f mice or AiCOP4 mice to express GCaMP6f or hChR2 in iSPNs. We imaged EYFP fluorescence in A2a-AiCOP4 mice and found high expression in the dorsal striatum, GPe, but not in the SNr (Fig. 1A). Using immunohistochemistry, we quantified the percentage of striatal neurons expressing GCaMP6f in the A2aCre-GCaMP6f mice. We labeled neurons using the pan-neuronal marker NeuN and GFP to visualize GCaMP6f-expressing cells and found that on average 42.9 + 4.9 % of striatal neurons expressed GFP (n = 3 mice, 5 micrographs/ mouse; n cells expressing GFP/ n of cells expressing NeuN in each mouse: 461/1021, 396/1064, 486/1050), indicating GCaMP expression in ~ half of the striatal SPNs (Fig. 1B) in accordance with previous reports (Graveland & Di Figlia, 1985). No GCaMP6f- expressing cell somata was found in the GPe, and sparse labeling was observed in the cortex. We functionally assessed the A2a-AiCOP4 line by performing whole-cell patch clamp recordings in the GPe (fig. 1C). We first assessed neurotransmission at iSPN-GPe synapses. We used a low-chloride intracellular and first recorded optically evoked postsynaptic currents while clamping the membrane potential near the sodium reversal potential (~0mV). The outward current recorded at this holding potential was decreased by Gabazine to 7 + 4.8 % of baseline (Fig. 1C) (n = 6 cells from n = 3 mice, paired t-test p < 0.0001). Next, we clamped the neurons near the chloride reversal potential (~ -45 mV) and recorded no current at this potential in 5/ 6 neurons. In 1/6 neurons, a DNQX and DL-AP5 insensitive, TTX sensitive current emerged. Taken together, these results indicate that optical stimulation of iSPN-GPe projections in the A2aCre-AiCOP4 line evokes GABA release. Next, we assessed whether the recorded oIPSCs were TTX- sensitive and monosynaptic (Fig. 1D). We found that TTX application abolished oIPSCs (amplitude decreased to 1 + 1.4 % of baseline), and oIPSC amplitude was restored to 195 + 60.9% of baseline by TTX + 4-AP co-application (n = 5 cells from n = 4 mice; mixed-effect analysis, F _(2, 11)_, treatment effect p < 0.0001; Tukey’s multiple comparisons test: baseline vs TTX p = 0.0021; Baseline vs TTX + 4- AP = 0.0023; TTX vs TTX + 4-AP p < 0.0001) (Fig. 1E). These results indicate that the oIPSCs were TTX-sensitive and monosynaptic, also according to their onset latency (Fig. 1F) (low-chloride, cells from 1D: 3.6 + 1.4 ms; high-chloride, cells from 1E: 1.2 + 0.37 ms) (Cho et al., 2013; Petreanu et al., 2009). We next tested whether the PreCaTs evoked in the GPe through local electrical stimulation (Fig. 1G) were activity- dependent and Ca^2+^ dependent. In the first experiment, we delivered stimuli of increasing intensity using 100 μA increments (range 0 μA- 700 μA) and found that the PreCaTs amplitude increased with stimulation intensity, indicating its activity-dependency (Fig. 1H) (n = 5 slices from n = 4 mice; one-way rmANOVA, F _(7,28)_ = 26.7, p < 0.0001). In the second experiment, we followed the PreCaTs amplitude during a baseline period of 10 minutes, and next we switched the recording solution from normal ACSF to Ca^2+^-free ACSF. The PreCaT amplitude decreased to 29.1 + 7 % of baseline during Ca^2+^- free ACSF application, and re-application of normal ACSF partially restored the PreCaTs to 50.8 + 12.5 % of baseline (Fig. 1I, J) (n = 5 slices, n = 4 mice one-way rmANOVA, Treatment: F _(2,8)_ = 71.94; p < 0.0001; Tukey’s multiple comparisons test: Ca^2+^ free vs baseline p < 0.0001; washout vs baseline p < 0.0001; washout vs Ca^2+^ free p = 0.0186). In summary, these validation steps indicated that the A2aCre-GCaMP6f and A2aCre-AiCOP4 lines represent a bona fide approach to study presynaptic calcium and GABA release from iSPN to GPe neurons.

### P/Q-type are the predominant VGCC controlling presynaptic Ca^2+^ and GABAergic transmission at the indirect pathway projections to the GPe

We performed a series of experiments to characterize the VGCCs mediating PreCaTs and GABAergic transmission at iSPN-GPe projections. We studied the effects of the bath application of selective blockers of P/Q type, N-type or L-type VGCCs on PreCaTs (Fig. 2A, B). Application of the P/Q-type VGCC blocker, ω-agatoxin IVA (ATX, 200 nM), decreased PreCaTs amplitude to 71.7 + 12.7 % of baseline and 53.3 + 18.5 % of baseline after washout (n = 6 slices from n = 6 mice, one-way rmANOVA, Treatment: F _(2, 10)_ = 32.53; p < 0.0001; Tukey’s multiple comparisons test: Baseline vs ATX p = 0.0017; Baseline VS Washout p < 0.0001; ATX VS Washout p = 0.0266). Application of the N-type VGCC blocker, ω-Conotoxin GVIA (CTX, 1 μM), did not significantly decrease in PreCaTs amplitude, which was 85.2 + 11.3 % of baseline after drug and 78 + 17.3 % of baseline following washout (n = 8 slices from n = 8 mice, one-way rmANOVA, Treatment: F _(2, 14)_ = 8.38; p = 0.004; Tukey’s multiple comparisons test: Baseline vs CTX p = 0.0565; Baseline VS Washout p = 0.0032; CTX VS Washout p = 0.322). Bath application of the L-type VGCC blocker, Nifedipine (10 μM), did not decrease PreCaTs amplitude at iSPN-GPe projections, which was 95 + 5.6 % of baseline and 90.2 + 7.8 % after washout (n= 5 slices from n = 5 mice, one-way rmANOVA, Treatment: _F (2, 8)_ = 3.638, P = 0.0752). Two-way rmANOVA comparison of ATX, CTX and Nifedipine application effects on PreCaTs at iSPN-GPe projections revealed a significant interaction between the time (timepoints analyzed: baseline, drug, washout) and drug factors, indicating a difference in the effect of the 3 blockers on PreCaTs (Time X drug: F _(4, 32)_ = 6.933; p = 0.0004; agatoxin vs nifedipine p = 0.004). Overall, these results indicate that P/Q type VGCCs are the primary controllers of PreCaTs at iSPN-GPe terminals. Next, we examined the effects of the bath application of selective blockers of P/Q type, N-type or L-type VGCCs on GABAergic transmission (Fig. 2C, D). Application of ATX induced a decrease in oIPSC amplitude to 20.2 + 14.4 % of baseline which persisted at 8.6 + 7.2 % of baseline after washout (iSPN-GPe, one-way rmANOVA, n = 5 cells from n = 4 mice, Treatment: F _(2, 8)_ = 189.1; p < 0 .0001; Tukey’s multiple comparisons test: Baseline vs ATX p < 0.0001; Baseline vs Washout p < 0.0001; ATX VS Washout p = 0.1172). Application of CTX induced a significant decrease in oIPSC amplitude to 73.2 + 12.7% of baseline which persisted at 69 + 22.6 % of baseline after washout (n = 5 cells from n = 4 mice, one-way rmANOVA, Treatment: F _(2, 8)_ = 8.4; p = 0.0108; Tukey’s multiple comparisons test: Baseline VS CTX p = 0.0275; Baseline VS Washout: p = 0.0133; CTX VS Washout p =0.8689). Application of Nifedipine induced a significant decrease in oIPSC amplitude to 78.9 + 11.1 % of baseline which persisted at 80.6 + 12.9 % of baseline after washout (n= 5 cells from n = 4 mice, one-way rmANOVA, Treatment: F _(2, 8)_ = 10.97, P = 0.0051; Tukey’s multiple comparisons test: Baseline VS Nifedipine p = 0.0074; Baseline VS Washout p = 0.0115; Nifedipine VS Washout p = 0.9437). Two-way RM ANOVA comparison of ATX, CTX and Nifedipine application effects on oIPSCs at iSPN-GPe synapses revealed a significant interaction between the time (timepoints analyzed: baseline, drug washout) and drug factors (Time X drug: F _(4, 24)_ = 21.09; p < 0.0001; Tukey’s multiple comparisons test: agatoxin vs conotoxin p < 0.0001; agatoxin vs nifedipine p < 0.0001).

**Figure 2:**
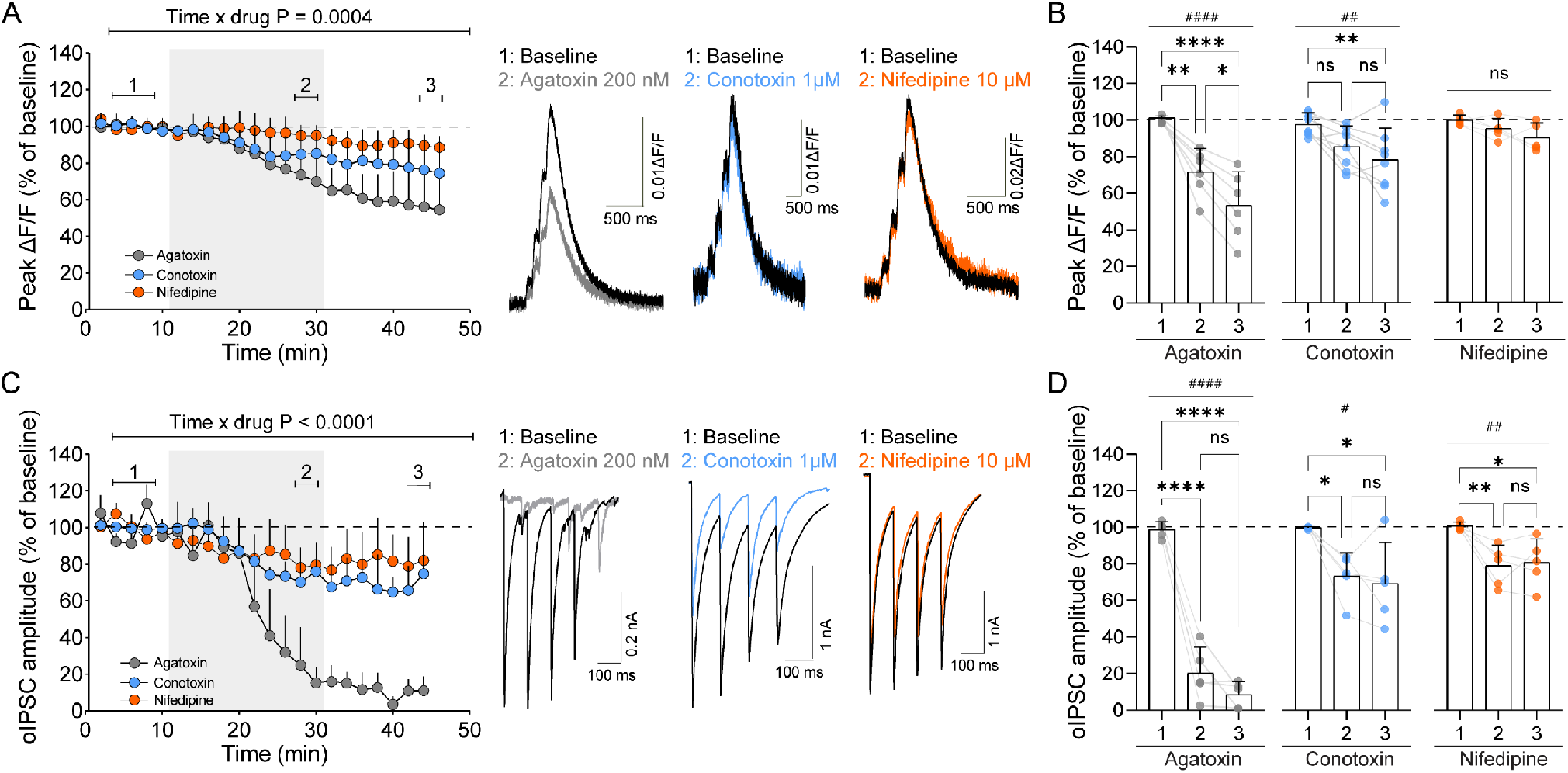
P/Q-type are the predominant VGCC controlling presynaptic Ca^2+^ and GABAergic transmission at the indirect pathway projections to the GPe. A: Timecourse of VGCC blocker application effects on PreCaTs and raw photometry traces. B: Summary of drug effects: baseline, last 2 minutes of drug, last 2 minutes of washout. Blockade of P/Q -type VGGCs (Agatoxin, n = 6 slices from n = 6 mice) significantly decreased PreCaTs from baseline, whereas no significant effect on PreCaTs was observed by blocking N-type VGCCs (Conotoxin, n= 8 slices from n = 8 mice) or L-type VGCCs (Nifedipine, n = 5 slices from n = 5 mice). C: Timecourse of VGCC blocker application effects on oIPSCs and raw electrophysiological traces. D: Comparison of drug effects: baseline, last 2 minutes of drug, last 2 minutes of washout. Blockade of P/Q -type VGGCs (Agatoxin, n = 5 cells from n = 4 mice) significantly decreased PreCaTs from baseline, and smaller but significant effects were induced by blocking N-type VGCCs (Conotoxin, n = 5 cells from n = 4 mice) or L-type VGCCs (Nifedipine, n = 5 cells from n = 4 mice). * p < 0.05; **/ ## P < 0.01; *** P < 0.0001; ****/ *####* P < 0.0001.

Taken together, these results indicate that P/Q type VGCCs are the primary VGCC involved in GABAergic transmission at iSPN-GPe synapses. In contrast, N-type and L-type VGCCs play a significant but smaller role.

### VGCC-dependent modulation of iSPN-GPe projections by GABA_B_ receptors

We studied the effects of GABA_B_ receptor activation on PreCaTs at iSPN-GPe projections (Fig. 3A, B). Bath application of the GABA_B_ agonist Baclofen (10 μM) decreased PreCaTs amplitude to 45.7 + 15.1 % of baseline which persisted at 50.8 + 23.3 % of baseline after washout (n = 7 slices from n = 5 mice, one-way rmANOVA, Treatment: F _(2, 12)_ = 40.94; p < 0.0001; Tukey’s multiple comparisons test: Baseline VS Baclofen p < 0.0001; Baseline VS Washout p < 0.0001; Washout VS Baclofen p = 0.7323). To test whether Baclofen application induced a long-term depression of PreCaTs at iSPN-GPe projections we performed a second experiment of pulse-chase where Baclofen application was followed by the application of the GABA_B_ antagonist CGP 35358 (2 μM) (Fig. 3C, D). Bath application of Baclofen decreased PreCaTs amplitude to 46.5 + 12.7 % of baseline, an effect that was reversed by CGP 35358 which brought PreCaTs to 107.5 + 23 % of baseline (n = 5 slices from n = 2 mice, one-way rmANOVA, Treatment: F _(2, 8)_ = 23.10; p = 0.0005; Baseline vs Baclofen p = 0.0014; Baseline VS CGP p = 0.7848; Baclofen VS CGP p = 0.0007).

**Figure 3:**
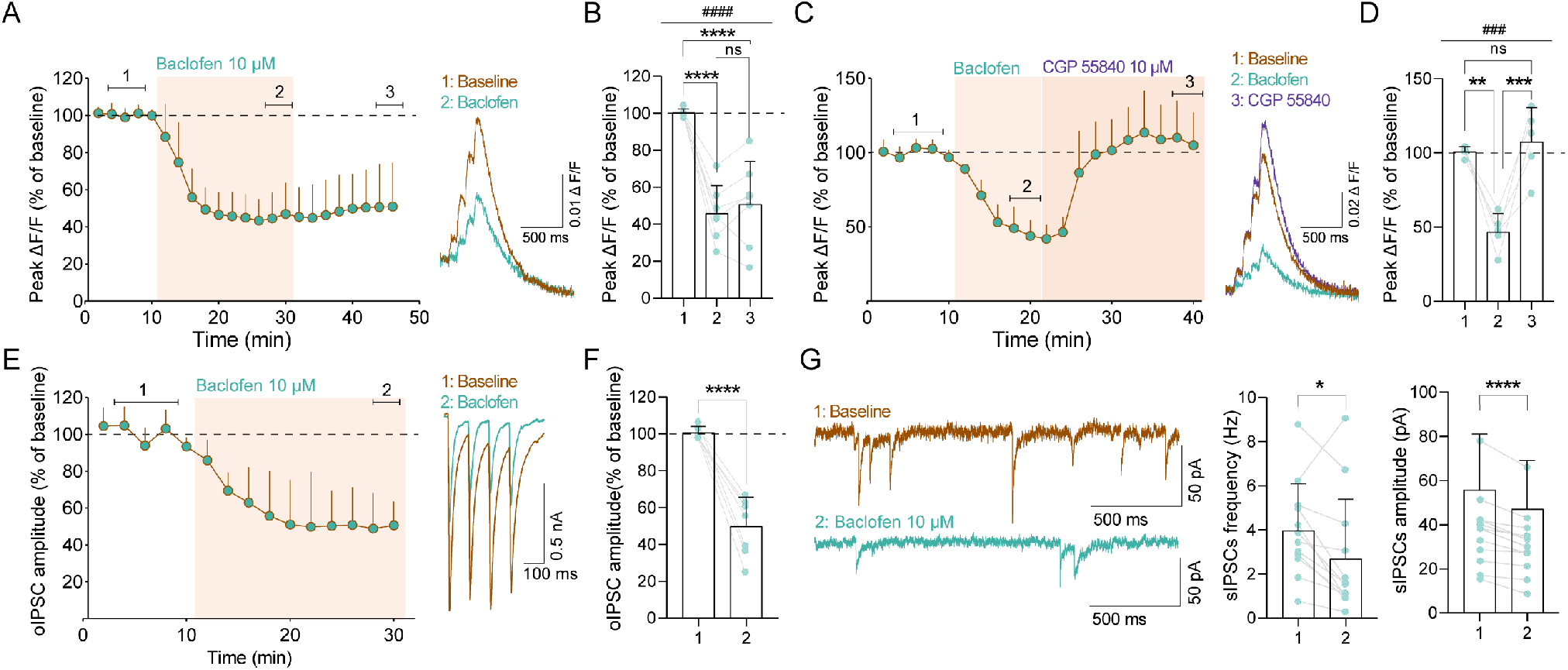
VGCC-dependent modulation of iSPN-GPe projections by GABA_B_ receptors. **A:** Time course of Baclofen effects on PreCaTs and raw photometry traces. **B:** Summary of drug effects: baseline, last 2 minutes of drug, last 2 minutes of washout. GABA_B_ activation (Baclofen, n = 7 slices from n = 5 mice) significantly decreased PreCaTs amplitude. **C:** Time course experiment of Baclofen followed by CBP 55840 effects on PreCaTs **D:** Baclofen significantly decreased PreCaTs and CGP55840 application reversed PreCaTs to baseline levels (n = 5 slices from n = 2 mice). **E:** Time course of Baclofen effects on oIPSCs at iSPN-GPe synapses and raw electrophysiological traces. **F:** Baclofen significantly reduced oIPSCs at iSPN-GPe terminals (n = 7 cells from n = 5 mice). **G:** Effects of Baclofen application on sIPSCs. Example traces and summary bar graphs. Baclofen application significantly decreased the frequency and amplitude of sIPSCs recorded in the GPe, indicating a presynaptic site of action (n= 12 cells from n = 10 mice). * p < 0.05; **/ ## P < 0.001; ***/ ### P < 0.001; ###/ **** P < 0.0001.

**Figure 4:**
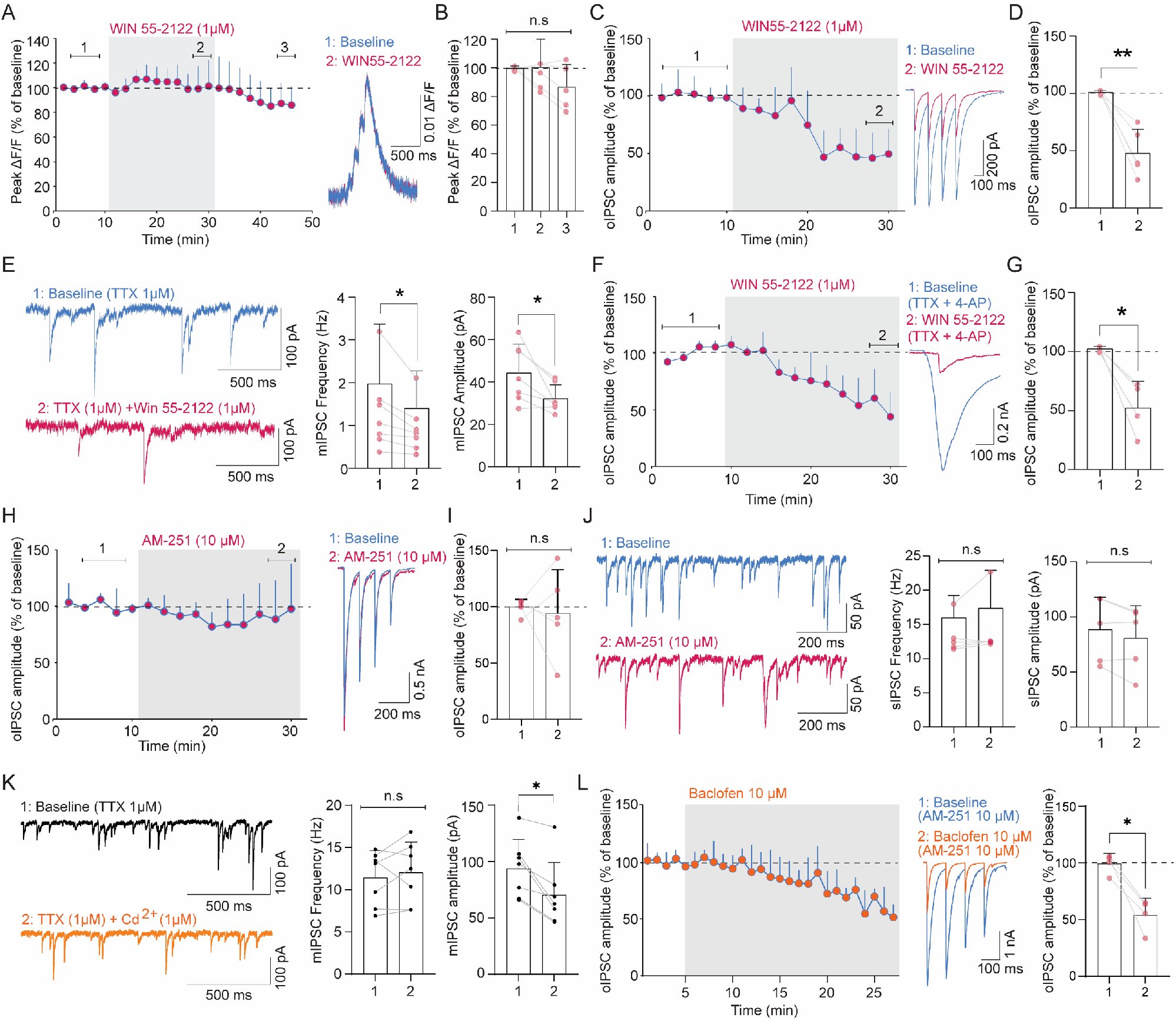
VGCC- and K_V_1-independent mechanisms of presynaptic modulation of iSPN-GPe projections by CB1 receptors. Time course of WIN effects on PreCaTs and raw photometry traces. **B:** Summary of drug effects: baseline, last 2 minutes of drug, last 2 minutes of washout. CB1 receptor activation (WIN 55-2122, n = 5 slices from n = 4 mice) did not change PreCaTs amplitude. **C:** Time course of WIN effects on oIPSCs at iSPN-GPe synapses and raw electrophysiological traces). **D:** WIN 55-2122 application significantly decreased oIPSC amplitude (n = 5 cells from n = 5 mice). **E:** Effects of WIN 55-2122 application on mIPSCs. Example traces and summary bar graphs. WIN 55-2122 application significantly decreased the frequency and amplitude of mIPSCs (n= 7 cells from n = 4 mice) **F**: Time course of WIN 55-2122 effects on oIPSCs recorded in the presence of TTX + 4AP. **G**: WIN 55-2122 significantly reduced oIPSCs at iSPN-GPe terminals (n = 4 cells from n = 3 mice). **H, I:** No effect of AM-251 application on oIPSCs (n = 5 cells from n = 3 mice). **J**: No effect of AM-251 application on sIPSC frequency or amplitude (n= 5 cells from n = 3 mice). **K**: Cd^2+^ decreased the amplitude but not frequency of mIPSCs (n = 7 cells from n = 3 mice). **L**: Baclofen effects were preserved in slices pre-incubated with AM-251 (n = 4 cells from n = 2 mice). * p < 0.05; ** P < 0.011.

We next examined the effects of GABA_B_ receptor activation on oIPSCs at iSPN-GPe projections (Fig. 3E, F). Bath application of the GABA_B_ agonist Baclofen decreased the olPSC amplitude to 49.8 + 16 % of baseline (n = 7 cells from n = 5 mice, paired t-test, p = 0.0002). To determine the site of action of GABA_B_ receptors in the GPe we analyzed sIPSCs (Fig. 3G). We found that Baclofen application induced a significant decrease in sIPSC frequency from baseline 4 + 2.1 Hz to Baclofen 2.3 + 2.7 Hz and amplitude from baseline 55.7 + 25.4 pA to Baclofen 47 + 22 pA (n = 12 cells from n = 10 mice; Frequency: paired t-test, p = 0.0195; Amplitude: paired t-test, p < 0.0001). The observed sIPSC decrease induced by baclofen application is indicative of a presynaptic effect, whereas a change in amplitude might reflect an effect on multivesicular release or a shift in the population of synapses contributing to the sIPSCs. Taken together, these experiments indicate that GABA_B_ receptor modulation of iSPN-GPe projections is reversible VGCC- mediated and involves presynaptic modulation of GABA release.

### VGCC- and K_v_1- independent mechanisms of presynaptic modulation of iSPN-GPe projections by CB1 receptors

We next studied the effects of CB1 receptor activation on PreCaTs at iSPN-GPe projections (Fig. 4 A, B). Bath application of the CB1 agonist WIN 55212-2 did not change PreCaTs amplitude, which was 100.3 + 19.7 % of baseline following WIN 55212-2 application and 86.6 + 15.7 % of baseline after washout (n = 5 slices from n = 4 mice, one-way rmANOVA, Treatment: F _(2, 8)_ = 2.386; p = 0.154). We next asked whether CB1 receptors reduced GABAergic transmission specifically at iSPN-GPe synapses (Fig. 4C, D). Bath application of WIN 55212-2 decreased the oIPSC amplitude to 47.8 + 20.8 % of baseline (n = 5 cells from n = 5 mice, paired t-test, p = 0.0053). CB1 activation did not decrease PreCaTs while inhibiting GABAergic transmission at iSPN-GPe projections. Next, we recorded mIPSCs in presence of the Na_v_ blocker TTX (Fig. 4E). Bath application of WIN 55212-2 decreased the mIPSC frequency from 2 + 1.4 Hz to 1.4 + 0.88 Hz (n = 7 cells from n = 4 mice, paired t-test, p = 0.0314) and decreased the mIPSC amplitude from 44.3 + 13.6 pA to 31.1 + 6.5 pA (n = 7 cells from n = 4 mice, paired t-test, p = 0.025). This indicates a presynaptic effect of CB1 receptor activation on GABA release in the GPe, but a change in amplitude might be indicative of an additional postsynaptic contribution, an effect on multivesicular release or a shift in the population of synapses contributing to the mIPSCs. We tested whether CB1 receptor effects at iSPN-GPe synapses might be mediated by K_V_1 channels, as suggested for glutamatergic synapses in the nucleus accumbens (Robbe et al., 2001) and cerebellar parallel fibers (Daniel et al., 2004). We examined WIN 55-2122 effects by recording baseline and drug application in aCSF containing the K_v_1 channel blocker 4-AP and TTX. Under these experimental conditions hChR2-evoked presynaptic depolarization is independent of Na_v_ channels and K_v_ channels, and monosynaptic connectivity is isolated (Cho et al., 2013; Petreanu et al., 2009). We found that WIN 55212-2 application produced a reduction of oIPSC amplitude to 52.3 + 22. 5 % of baseline (n = 4 cells from n = 3 mice, paired t-test, p = 0.0216) (Fig. 4F; G). These experiments indicate that CB1 effects on iSPN-GPe projections are monosynaptic and not mediated through the inhibition of K_V_1 or Na_V_ channels. To test whether an endocannabinoid tone is present on iSPN-GPe projections, we bath applied the CB1 antagonist AM-251. We found that AM-251 (10 μM) application did not significantly change the amplitude of oIPSCs (amplitude: 94.3 + 38.6 % of baseline after AM-251; paired t-test, p = 0.776), nor the amplitude and frequency of sIPSCs recorded in the GPe (Fig 4H-I-J) (n = 5 cells from n = 3 mice; frequency: baseline 16 + 3.2 Hz, AM-251 17.4 + 5.5 Hz; paired t-test, p = 0.2767; amplitude: baseline 88.5 + 29.6 pA, AM-251 80.7 + 29.6 pA; paired t-test, p = 0.1159).

To test for a possible contribution of VGCCs to mIPSCs in the GPe, we studied the effects of the non-specific VGCC blocker Cd^2+^ on mIPSCs (Fig. 4K). We found that Cd^2+^ application did not change the frequency of mIPSCs in the GPe (n = 7 cells from n = 3 mice; baseline: 11.4 + 3.2 Hz; Cd^2+^: 12 + 3.6 Hz; paired t-test, p = 0.4728), ruling out presynaptic effects of Cd^2+^. Cd^2+^ application decreased the amplitude of mIPSCs in the GPe (n = 7 cells from n = 3 mice; baseline 93.7 + 26 pA; Cd^2+^ 70.3 + 28.9 pA; Wilcoxon signed-rank test, p = 0.0156), indicating a possible post-synaptic effect of Cd^2+^ mediated by direct Cd^2+^ modulation of GABA_A_ receptors (Molnár et al., 2004) or indirect by altered Ca^2+^- dependent modulation of GABA_A_ receptors (Grayelle et al., 2021). Taken together, Na_v_ independent GABA release in the GPe is independent from VGCCs.

Finally, we sought to establish if a cross- talk exist between GABA_B_ and CB1 receptors on iSPN-GPe projections. In a subset of cells from the experiments in Fig. 4H, AM-251 bath application was followed by the bath application of Baclofen. We found that bath application of Baclofen (10 μM) in presence of AM-251 (10 μM) significantly decreased the oIPSC amplitude to 54.19 + 14.6 % of baseline (Fig. 4L) (n = 4 cells from n = 2 mice; paired t-test, p = 0.0149). Taken together, these results indicate that CB1 receptors control iSPN-GPe projections through a mechanism independent from VGCCs, K_V_1 or Na_V_ channels. Furthermore, we found no evidence for cross-talk between GABA_B_ and CB1 receptor mediated control of GABA release on iSPN-GPe projections.

## Discussion

Here we used a combination of brain slice photometry and whole-cell patch clamp to study the presynaptic modulation of indirect pathway projections (iSPN-GPe) by CB1 and GABA_B_ receptors. We first determined that Presynaptic Calcium Transients (PreCaTs) and GABA release at striatal iSPN-GPe terminals are primarily mediated by P/Q-type VGCCs. GABA_B_ receptors modulate PreCaTs and GABA release at these terminals through a presynaptic mechanism. CB1 receptors control GABA release but not PreCaTs through a VGCC-, Na_V_ and K_V_1 independent presynaptic mechanism. We found no evidence of cross- talk between GABA_B_ and CB1 receptors on iSPN-GPe projections. Our results indicate that GABA_B_ and CB1 receptors mediate presynaptic inhibition at iSPN-GPe synapses through distinct mechanisms.

We focused our experiments on the study of presynaptic modulation of iSPN-GPe projections assuming similar principles governing PreCaTs and GABA release at iSPN-GPe synapses on distinct GPe neuronal populations. However, presynaptic modulation of iSPN projections might be recruited with specific modalities by different neuronal populations of GPe neurons.

### Control of presynaptic Ca^2+^ at iSPN-GPe projections

We studied PreCaTs at iSPN-GPe projections using a genetic approach to express GCaMP6f in iSPNs. As a result, GCaMP6f was expressed in iSPNs independently from their location in striatal subregions (dorsomedial, ventromedial, or dorsolateral striatum) or the molecularly defined striatal subcompartments, patch and matrix. Using field photometry and pharmacology we were able to measure activity- dependent PreCaTs and determine which VGCCs subtypes mediate them. Further, we could study their modulation by GABA_B_ and CB1 receptors. Field photometry allowed the measurement of PreCaTs in a selected field of view within the GPe containing iSPN axons. This population-level measurement likely reflects the sum of axonal Ca^2+^ proximal to, and distal from, presynaptic terminals/ release sites given the widespread expression of the fluorescent indicator throughout the axon. A limitation of this approach is that the spatial resolution of field photometry might not be sufficient to reflect small Ca^2+^ variations restricted to the presynaptic terminals but critical for neurotransmitter release. To counter this limitation, we generated mice genetically expressing hChR2 in iSPNs and performed patch-clamp recordings to measure neurotransmitter release. This approach ensured high temporal and spatial resolution allowing us to examine monosynaptic iSPN-GPe projections.

The use of whole cell patch-clamp allowed us to study activity- dependent neurotransmitter release and its control by VGCCs, GABA_B_ and CB1 receptors. It also allowed to measure action potential-independent, VGCC- independent neurotransmitter release by recording mIPSCs in the presence of the Na_V_ blocker TTX. We found that mIPSC frequency was insensitive to the blockade of VGCCs by Cd^2+^ indicating that mIPSCs in the GPe are VGCC- independent. Taken together, a combination of field photometry, whole cell patch-clamp and optogenetics allowed us to gain mechanistic insights on the modulation of iSPN projections by GPCRs. In the present study we examined GPCR modulation using bath application of agonists. While this approach ensures uniform concentrations of agonist throughout the preparation, the drug application time and washout are much slower than with other techniques (e.g. rapid, local agonist application). In addition, while there is the possibility with this approach that agonist actions may occur at sites other than the presynaptic terminal of the synapse under study, this concern is mitigated to some extent by the examination of sIPSCs in the presence of TTX which will reduce many neuronal interactions that could give rise to indirect effects, as well as the pharmacological isolation of ISPCs that reduces influences from glutamatergic synapses.

We found that PreCaTs were inhibited by an antagonist of P/Q type VGCC, but not by antagonists of N-type or L-type VGCC. GABA release was decreased by antagonists of P/Q type and to a smaller extent by antagonists of N-type or L-type VGCCs. One possibility is that small variations in presynaptic Ca^2+^ that were not detectable using slice photometry contributed to the effects of N-type of L-type antagonists on GABA release.

### Control of iSPN-GPe projections by GABA_B_ receptors

We found that GABA_B_ receptors modulate the iSPN-GPe projection by suppressing GABA release and PreCaTs. GABA_B_ receptor activation decreased sIPSC frequency in the GPe, indicating a presynaptic site of action. A decrease in sIPSC frequency and amplitude does not exclude GABA_B_ receptor-mediated inhibition of other GABAergic inputs to the GPe, including local collaterals, bridge collateral dSPN projections and GPi inputs. A decrease in sIPSC amplitude might result from the inhibition of selected GABAergic inputs or from the reduction of multi-vesicular release. Alternatively, these changes may emerge from post-synaptic GABA_B_ mediated effects (Kaneda & Kita, 2005). Our results are consistent with previous reports indicating VGCCs as one of the downstream targets of GABA_B_ receptor signaling at corticostriatal synapses (Kupfershmidt & Lovinger, 2015) granule-Purkinje cell cerebellar synapses (Dittman & Regehr, 1996) and hippocampal synapses (Doze et al., 1995; Wu & Saggau, 1995). Ultrastructural studies have identified the expression of GABA_B_ receptors in the GPe in rodents and primates. Pre-synaptic GABA_B_ receptors were found on GABAergic inputs to the GPe and on anterogradely labelled striatopallidal projections (Charara et al., 1999; Chen et al., 2004). Further, GABA_B_ receptors were found post-synaptically at GABAergic synapses and presynaptically at glutamatergic synapses in the GPe (Chen et al., 2004).

Hence, GABA_B_ receptors are positioned to regulate GPe physiology as autoreceptors, heteroreceptors or postsynaptic receptors. In our recording conditions, no GABA_B_ mediated postsynaptic responses were observed since GIRK channels were blocked by including Cs^+^ in the intracellular recording solution.

### Control of iSPN-GPe projections by CB1 receptors

Our results indicate that CB1 receptors modulate iSPN-GPe projections by reducing GABA release while our slice photometry experiments indicated no effect of CB1 receptor activation on presynaptic Ca^2+^. Further, we found that bath application of the CB1 agonist WIN 55212-2 induced a decreased mIPSC frequency and reduced the amplitude of oIPSCs recorded in presence of K_v_1 and Na_v_ antagonists. Overall, our presynaptic Ca^2+^ measurements and neurotransmitter release measurements converged in indicating a VGCC-independent, Na_V_ and K_V_1 independent control of GABA release by CB1 receptors on iSPN projections to the GPe. These results reproduce, extend, and provide synapse specificity to previously published work on the inhibitory effect of CB1 receptors on GABAergic synaptic transmission in the GPe (Engler et al., 2006). Engler and colleagues reported a CB1-mediated inhibition of presynaptic calcium at striatopallidal synapses, while we observed no CB1- mediated inhibition of PreCaTs. These discrepant results might be due to differences in the slice preparation used, differences between mouse strains, and differences in analysis methods used to determine PreCaTs. We found no evidence for a CB1- mediated tonic inhibition of GABA release on iSPN-GPe projections indicating that the lack of effect of the CB1 agonist WIN 55212-2 in our experiments is not ‘masked’ by a tone. CB1 receptors have been proposed to suppress neurotransmitter release through the activation of K_v_ channels at cortico-accumbal (Robbe et al., 2001) and cerebellar parallel fiber synapses (Daniel et al., 2004). CB1-mediated inhibition of Na_v_-independent GABA release in the DS was also observed (Adermark et al., 2009). Other studies proposed that presynaptic CB1 receptors depress glutamate release at corticostriatal through the inhibition of N-type Ca^2+^ channels (Huang et al., 2001), and a similar VGCC-dependent action was reported at hippocampal GABAergic synapses (Hoffman & Lupica, 2001). CB1 receptors were also shown to depress presynaptic Ca^2+^ influx at cerebellar granule-Purkinje cell synapses (Brown et al., 2004), as well as Ca^2+^ currents at calyx of Held synapses in the auditory brainstem (Kushmerick et al., 2004). The signaling cascade recruited by CB1 receptor activation on iSPN-GPe terminals remains to be fully elucidated. A possible alternative mechanism might involve the direct interaction of the G_Bγ_ subunit with SNARE complexes. It has been proposed that G_Bγ_ subunits can target the C-terminus domain of SNAP-25 (Zurawski et al., 2019). This mechanism has been shown to mediate effects of the 5HT1B receptor in the hippocampus, and the α2a adrenergic receptor on glutamatergic synapses in the BNST, (Zurawski et al., 2019), as well as effects of GABA_B_ receptors on glutamatergic inputs to SPNs in the nucleus accumbens (Manz et al., 2019). However, CB1 receptors have not yet been shown to activate this signaling cascade. A different possibility comes from a study that examined CB1-mediated control of GABA release at dSPN-SNr projections (Soria-Gomez et al., 2021). The authors found that CB1 receptors reside on the plasma membrane and on mitochondria in this projection. Plasma membrane-associated CB1 receptors controlled substance P release via inhibition of cytosolic PKA, whereas mitochondria associated CB1 receptors controlled GABA release via inhibition of mitochondrial PKA.

Here, we found that CB1 signaling at iSPN-GPe synapses is non- VGCC dependent, fitting with a growing complexity of CB1 receptor mediated signaling in the CNS.

What is the functional significance of CB1 receptor signaling on iSPN-GPe terminals? The levels of the CB1 receptor in the GPe are among the highest in the CNS. CB1 receptors expressed in SPNs control locomotor and exploratory behavior (Bonm et al., 2021). The endocannabinoids anandamide and 2-arachidonoylglycerol (2-AG) were detected in the rat GPe (Di Marzo et al., 2000) but the mechanisms driving their release remains unclear. Local micronjections of synthetic cannabinoids in the GPe were found to increase ipsilateral rotations (Sañudo-Peña et al., 1999) and to enhance muscimol-induced catalepsy (Pertwee & Wickens, 1991). Further, reserpine treatment enhanced anandamide and 2-AG levels in the rat GPe (Di Marzo et al., 2000). CB1 receptors in the GPe might be primarily recruited by endocannabinoids released retrogradely. Alternatively, astrocytes surrounding striatopallidal projections (Galvan et al., 2010) might provide an alternative source of endocannabinoids, although these remain untested hypotheses. A third possibility involves endocannabinoid production and release at iSPN-GPe synapses recruited by neuromodulatory or glutamatergic inputs to the GPe. We did not find any evidence of endocannabinoid activation of CB1 receptors on striatopallidal neurons under the afferent stimulation or sIPSC recording conditions in this study. However, other patterns of afferent and neuronal activation need to be explored, perhaps with the help of newly developed endocannabinoid sensors (Dong et al., 2022).

We show that CB1 and GABA_B_ receptors recruit distinct presynaptic signaling mechanisms. By inhibiting VGCCs, GABA_B_ receptors might shunt action potential evoked neurotransmitter release at iSPN-GPe synapses, in response to increased presynaptic activity (autoreceptor role) or increased local GABA (heteroreceptor role). CB1 receptors might function synergistically with GABA_B_ receptors by further decreasing GABA release from striatopallidal synapses. We found that pre-incubation of the slices with a CB1 receptor antagonist did not prevent the effects of the GABA_B_ agonist, baclofen, indicating no cross talk between GABA_B_ and CB1 receptors at iSPN-GPe projections. The growing understanding of the GPe cell-type composition, connectivity, microcircuitry and function requires a synapse-specific approach to contextualize the role of CB1 and GABA_B_ receptor signaling in this region. A detailed analysis of the striato-pallidal and pallido-pallidal circuitry has proposed that iSPN-GPe projections are the densest in the GPe and primarily target prototypic over arkypallidal GPe neurons (Cui et al., 2021; Ketzef & Silberberg, 2021). PV+ neurons represent the primary source of collateral inhibition in the GPe (Cui et al., 2021; Higgs et al., 2021; Ketzef & Silberberg, 2021). Presynaptic inhibition of iSPN-GPe projections might therefore produce primarily a disinhibition of the output of GPe prototypic neurons to the SNr, Subthalamic Nucleus and Parafascicular Thalamus (Lilasharoen et al., 2020; Pamukcu et al., 2020) in addition to disinhibiting local collaterals. Presynaptic modulation will ultimately contribute to determine the GPe behavioral output, contributing to movement control and action selection.

## Conclusion

In conclusion, our results indicate that distinct mechanisms of presynaptic modulation are recruited by CB1 and GABA_B_ receptors on the striatal indirect pathway projections to the GPe. Presynaptic modulation by CB1 and GABA_B_ receptors decrease the synaptic efficacy of iSPN-GPe projections by inhibiting GABA release and ultimately contributes to shape the basal ganglia output. Our results encourage research on the synapse-specific mechanisms of Gi/o coupled receptor signaling which might inform the development of targeted pharmacological treatments for movement and psychiatric disorders.

## Author contributions

KPA, DL, GS and DML were responsible for the conception and design of the study. KPA, DL, GS performed electrophysiological recordings. GS analyzed electrophysiological recordings. KPA, DL and GMC performed slice photometry experiments and analyses. GS performed and analyzed immunohistochemical experiments. GS drafted the manuscript and the figures with inputs from all authors and all authors gave final approval.

## Acknowledgments

This work was supported by the National Institutes of Health, National Institute on Alcohol Abuse and Alcoholism, Division of Intramural Clinical and Biological Research (ZIAAA000416). We thank the NIAAA animal care stuff for their precious support for animal husbandry and veterinary care. We thank Guoxiang Luo for genotyping assistance, and all the Laboratory for Integrative Neurosciences for the helpful discussions.

## Data availability statement

All data generated or analyzed in this manuscript are included in the manuscript and its supplementary file(s).

## Notes

### Competing Interest Statement

The authors have declared no competing interest.

### Summary of Updates

Manuscript and figures revised. Data added.

